# Tumor microenvironment-induced FOXM1 regulates ovarian cancer stemness

**DOI:** 10.1101/2023.10.30.564779

**Authors:** Chiara Battistini, Hilary A. Kenny, Valentina Nieddu, Valentina Melocchi, Alessandra Decio, Alessia Gatto, Mariacristina Ghioni, Francesca M. Porta, Raffaella Giavazzi, Nicoletta Colombo, Fabrizio Bianchi, Ernst Lengyel, Ugo Cavallaro

## Abstract

In ovarian tumors, the omental microenvironment profoundly influences the behavior of cancer cells and sustains the acquisition of stem-like traits, with major impacts on tumor aggressiveness and relapse. Here, we exploit a patient-derived platform of organotypic cultures to study the crosstalk between the tumor microenvironment and ovarian cancer stem cells. The pro-tumorigenic transcription factor FOXM1 is specifically induced by the microenvironment in ovarian cancer stem cells, through activation of FAK/YAP signaling. The microenvironment-induced FOXM1 sustains stemness, and its inactivation reduces cancer stem cells survival in the omental niche and enhances their response to the PARP inhibitor Olaparib. By unveiling the novel role of FOXM1 in ovarian cancer stemness, our findings highlight patient-derived organotypic co-cultures as a powerful tool to capture clinically relevant mechanisms of the microenvironment/stem cells crosstalk, contributing to the identification of tumor vulnerabilities.

## Introduction

Epithelial ovarian cancer (OC) is the most lethal gynaecological cancer worldwide ^1^, and high-grade serous OC (HGSOC) is its most frequent and aggressive subtype, with a survival rate of only 30% at 5 years ^2^. HGSOC is often diagnosed at an advanced stage, when the disease has already spread into the peritoneal cavity ^3^. Moreover, despite an initial response to first-line treatments, 70% of HGSOC relapse within 2 years ^1^, and almost all recurrent HGSOC ultimately develop chemoresistance and become unresponsive to standard treatments ^4^.

Among the factors implicated in the aggressiveness of HGSOC, a role has been attributed to a small subpopulation of cells named ovarian cancer stem cells (OCSC). These cells have the ability to self-renew and to initiate tumorigenesis, therefore they are crucially involved in the process of peritoneal dissemination and colonization; moreover, they are intrinsically resistant to cytotoxic treatments, thereby evading standard chemotherapy and fueling tumor re-growth and relapse ^4^. These features make OCSC an optimal therapeutic target to tackle OC, but to reach this goal there is an urgent need to gain further insights into OCSC biology. In this context, the tumor microenvironment (TME) has been implicated in OC stemness, thus promoting the acquisition of an aggressive phenotype ^5^. Moreover, previous studies have demonstrated that the TME impacts significantly the response of OC cells to chemotherapy ^6^. Therefore, it is fundamental to generate preclinical *in vitro* models that incorporate the TME, to better mimic the OC biology and predict more faithfully its response to therapies, and, importantly, to address the specific mechanisms of its crosstalk with OCSC.

The peritoneal surface is the preferential site of HGSOC dissemination and offers an optimal niche to metastasizing cells ^7^. To recapitulate this process *in vitro*, three-dimensional multicellular organotypic models have been established using human omental fibroblasts and mesothelial cells. Adding OC cells to organotypic co-cultures has allowed to elucidate some of the molecular mechanisms that govern the OC/TME crosstalk ^8–11^. However, these studies have relied on established OC cell lines, which have an intrinsically limited disease relevance. Furthermore, while the organotypic setting appears an optimal tool to explore the influence of peritoneal TME on OC stemness-related features, so far it has not been applied to this objective.

In the current study, we report the discovery of FOXM1 as a major effector of the TME/OCSC interplay. FOXM1 is a transcription factor belonging to the conserved forkhead box (FOX) family, which has been associated with poor prognosis in several cancer types. The TCGA molecular profiling of OC reported altered FOXM1 signaling in 84% of cases ^12^, in agreement with the functional contribution of the transcription factor to various aspects of OC malignancy ^13^. Yet, whether FOXM1 is involved in OC stemness and in the pathophysiology of OCSC has remained elusive.

Here, we exploited a platform of clinically relevant patient-derived organotypic co-cultures to shed light on the crosstalk between OCSC and their TME. Notably, the TME-mediated induction of FOXM1 in OCSC is crucial for their survival in the omental niche, but also a liability that, if successfully targeted, may improve the response of tumor cells to drug treatments.

## Results

### Contact with TME induces transcriptional reprogramming in bulk OC cells and OCSC

To investigate the crosstalk between OC cells and their microenvironment in a clinically relevant setting, we reconstructed *in vitro* a 3D organotypic model of ovarian TME, entirely composed of patient-derived primary cells. We used tumor cells derived from ascites of HGSOC patients, cultured either as bulk, adherent population or as clonal spheres under non-adherent conditions. This method exploits fundamental properties of CSC, namely their ability to resist *anoikis*, to self-renew, and to proliferate when seeded at low density in non-adherent conditions, to enrich primary tumor cultures for cells endowed with stem-like features ^4^. The peritoneal microenvironment was recapitulated by the co-culture of primary mesothelial cells and fibroblasts derived from human omental biopsies ^14^. Single-cell suspensions of primary OC cells, cultured either as adherent cells (referred to as “bulk”) or as OCSC-enriched spheres, were labeled with CMFDA (a green fluorescent dye) and then seeded on top of omental stromal cells, to generate 3D organotypic cultures of ovarian TME (Fig. 1A).

**Figure 1.**
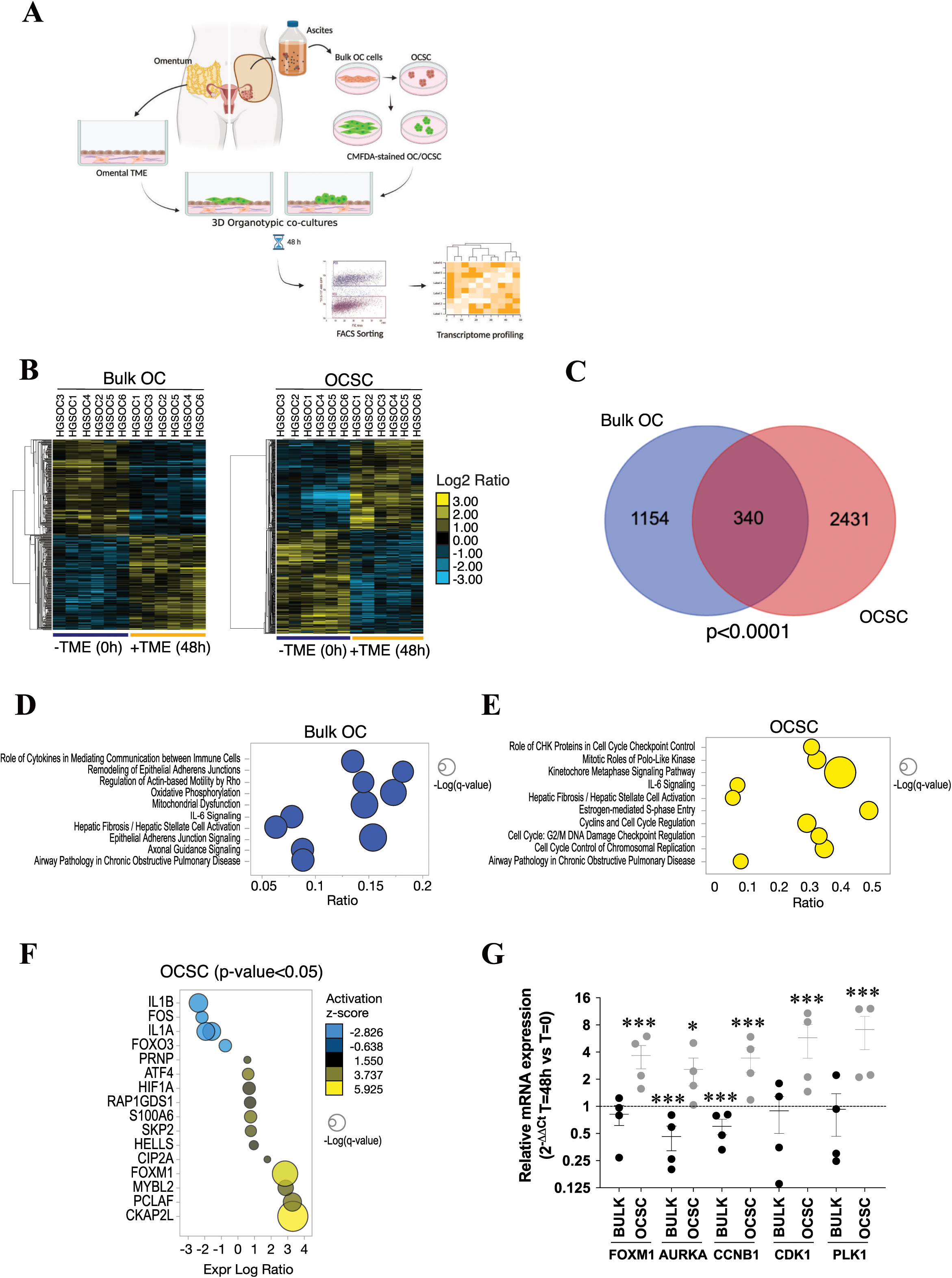
The TME induces transcriptional reprogramming in bulk OC cells and OCSC and activates FOXM1 pathway in OCSC. (A) Schematic representation of the patient-derived organotypic model. CMFDA-labelled bulk OC cells and OCSC are FACS-isolated after 48 hours of co-culture with TME (fibroblasts and mesothelial cells) and subjected to RNA sequencing. Created with BioRender.com. (B) Hierarchical clustering analysis of differentially expressed genes in bulk OC and OCSC of 6 ovarian cancer samples (HGSOC1-6) with or without the interaction with tumor microenvironment (TME). The heatmap shows gene expression change (log_2_ ratios) as per the legend. (C) Venn diagrams showing the number of DEGs regulated by TME in bulk OC and OCSC (hypergeometric test *p* = 2.377e-19). DEGs are listed in Supplementary Tables S2 and S3. (D, E) Bubble plots showing results of the Canonical Pathways Analysis of Ingenuity Pathways Analysis (IPA) in bulk OC (D) and OCSC (E). Bubble size represents the inverse of Logarithmic (-Log) of significance (q-value); for each pathway, ratio is the number of genes in our list over the total number of genes in the specific pathway. Only pathways with q-value ≤0.05 are shown. (F) Bubble plot showing the results of Upstream Regulator Analysis of IPA. Upstream modulators are predicted to be modulated by contact with TME in OCSC and are listed in Supplementary Table S4. Bubble size is proportional to inverse of Logarithmic (-Log) of significance [e.g., -Log(p-value), p-value <0.05; student’s t-test] of z-score. Bubble colors refer to activation z-score, as per the legend. Only modulators with coherent expression trend with the activation z-score are shown. (G) mRNA expression of *FOXM1* and its target genes (*AURKA*, Aurora Kinase A, *CCNB1*, Cyclin B1, *CDK1*, Cyclin-dependent Kinase 1, *PLK1*, Polo-like Kinase 1) was analyzed by qRT-PCR in 4 of the 6 patients profiled in panel B (HGSOC1-4). Data are represented as a relative mRNA expression (2^-ΔΔCt^) of bulk OC cells (in black) and OCSC (in grey) co-cultured with TME for 48 hours (T=48h) compared to cells grown in the absence of TME (T=0, dashed line). The dots represent each gene’s relative mRNA expression in each sample, while bars and whiskers represent its mean ± SEM among the 4 samples analyzed. Comparisons between experimental groups were done with two-sided Student’s t-test; \**p* < 0.05, \*\*\**p* < 0.005.

After 48 hours, 3D co-cultures were dissociated to single cells and CMFDA-labeled HGSOC cells were isolated by FACS and subjected to RNA-seq. Six independent 3D organotypic co-cultures (i.e., from six different patients, Suppl. Table S1) were analyzed and tumor cells from the co-cultures were compared to cells that have not been in contact with TME.

Overall, peritoneal TME induced a dramatic transcriptional reprogramming in HGSOC cells, with 1494 differentially expressed genes (DEGs) in bulk cancer cells and 2771 DEGs in OCSC (Fig. 1B and Suppl. Tables S2 and S3). A large fraction of DEGs were specific to either bulk OC or OCSC (Fig. 1C; p<0.001, hypergeometric test), indicating that the TME affected the two cell subpopulations in a different manner. Interestingly, Ingenuity Pathway Analysis (IPA) showed that the top regulated pathways in DEGs specific to bulk OC were mainly related to cytoskeletal remodeling, Rho-mediated motility, and metabolic processes such as oxidative phosphorylation (Fig. 1D). In contrast, pathways up-regulated in OCSC when co-cultured with the TME related to cell cycle regulation, S-phase entry and chromosomal replication (Fig. 1E).

These data demonstrated that TME exerted a distinct transcriptional program in primary OC cells and OCSC, yet with a remarkably divergent response in the two cell populations.

### TME induces FOXM1, which is required for OC stemness

We performed an Upstream Regulator Analysis by IPA to identify molecular hubs regulated by peritoneal TME in OCSC. One of the most prominent transcriptional regulators activated upon contact with the TME was FOXM1 (z-score=5.088; p-value= 2.61*10-20; Fig. 1F; Suppl. Table S4), a major driver of proliferation, survival and tumorigenesis ^15^. Gene Set Enrichment Analysis (GSEA) indeed showed a significant enrichment of the FOXM1 pathway in OCSC when cultured on TME (NES=+2.29; q-value<0.001) Suppl. Fig. 1A). Next, we performed qPCR analysis of FOXM1 expression, which confirmed the TME-induced activation of FOXM1 (Fig. 1G); qPCR also showed a set of cell cycle-related FOXM1 target genes (*AURKA*, *CCNB1*, *CDK1,* and *PLK1*) to be induced in OCSC cultured on TME (Fig. 1G). Moreover, immunofluorescence staining showed the presence of FOXM1 in the nuclei of primary, HGSOC-derived OCSC cultured on TME, and not in the same OCSC in the absence of TME (Suppl. Fig. 1B).

Notably, when we interrogated by GSEA a large cohort of HGSOC patients, i.e. the TCGA-OC dataset (N=535; see methods), we observed the enrichment for pathways related to cell cycle progression in high-FOXM1 compared to low-FOXM1 tumors (Suppl. Fig. 2), in line with the effect of TME on OCSC. This supported the ability of our organotypic model to recapitulate clinically relevant, FOXM1-mediated traits of the disease. In this context, FOXM1 expression was widespread across all major OC subtypes (Fig. 2A), supporting its relevance in ovarian tumorigenesis.

**Figure 2.**
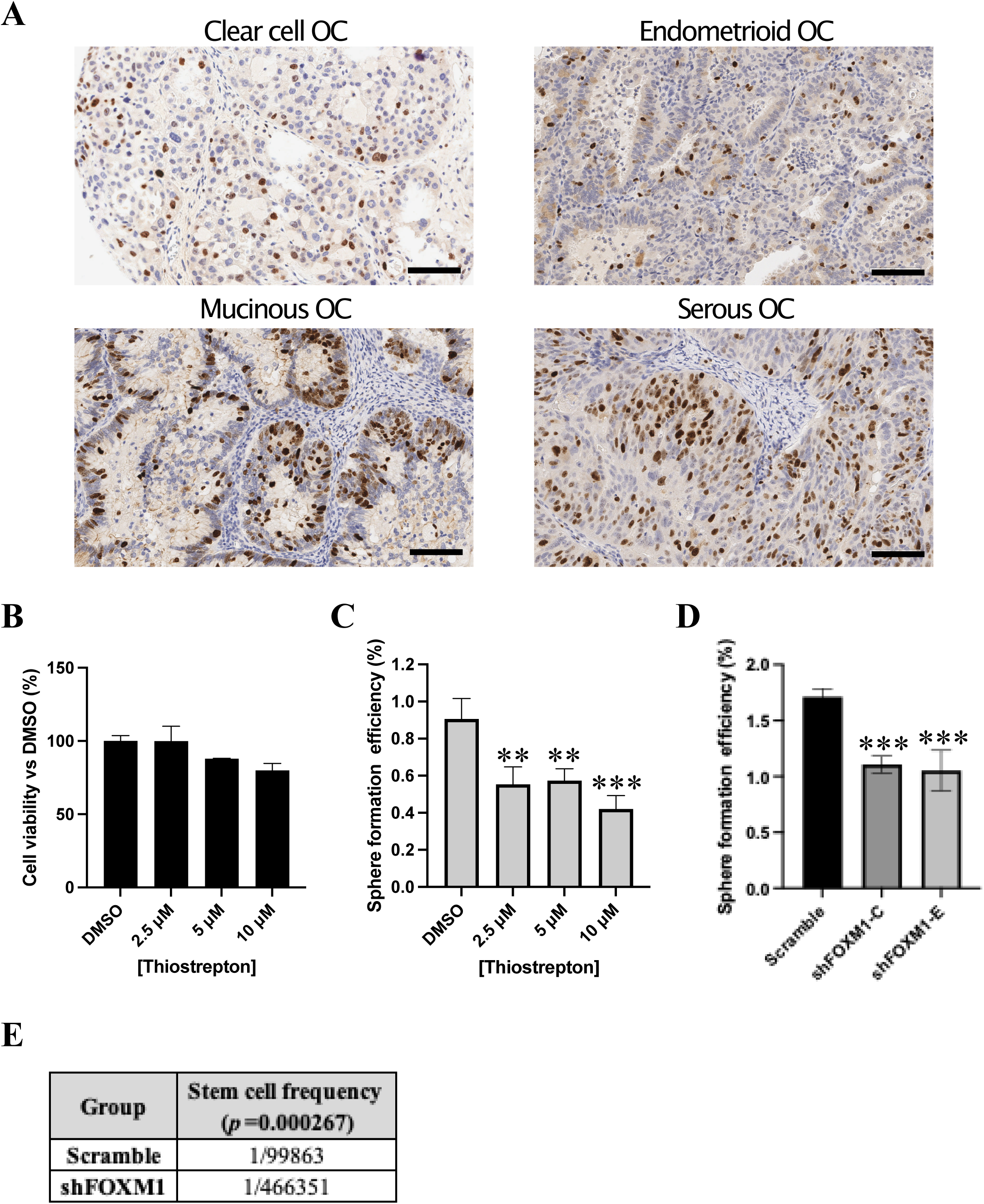
FOXM1 sustains stemness in OC. (A) IHC staining for FOXM1 on sections from different subtypes of OC. Sections were counterstained with Hematoxylin. Scale bars: 100 μm. (B) Viability of HGSOC1 primary cells measured after 72h of treatment with vehicle (DMSO) or 3 different doses of FOXM1 inhibitor Thiostrepton. (C) SFE in HGSOC1 primary sample treated with vehicle (DMSO) or with 3 different doses of FOXM1 inhibitor Thiostrepton. Comparisons between experimental groups were done with two-sided Student’s t-test; \*\**p* < 0.01, \*\*\**p* < 0.005. (D) Sphere formation assay performed on TYK-nu cells expressing either a control shRNA (Scramble) or 2 different shRNAs against FOXM1 (shFOXM1-C and E). Comparisons between experimental groups were done with two-sided Student’s t-test; \*\*\**p* < 0.005. (E) Nude mice were transplanted subcutaneously with decreasing numbers of either TYK-nu Scramble or TYK-nu shFOXM1-C cells, tumor take was assessed 21 days after injection (see Suppl. Fig. 3E) and an extreme limiting dilution assay (ELDA) was carried out to calculate the stem cell frequency (*p*=0.000267).

The role of FOXM1 in OC stemness has been studied only in cell lines and with limited mechanistic insights ^16–18^. Thus, we initially explored the functional contribution of FOXM1 to stemness in clinically relevant models using primary HGSOC cultures and blocking FOXM1 function with the inhibitor Thiostrepton, a natural cyclic oligopeptide antibiotic ^19^. Thiostrepton had no significant effect on the viability of bulk, adherent OC cells (Fig. 2B and Suppl. Fig. 3A), while it efficiently inhibited sphere formation (Fig. 2C and Suppl. Fig. 3B), suggesting the specific involvement of FOXM1 in promoting stem-like properties.

To gain further insights into the role of FOXM1 in OC stemness, we sought to identify an experimental model suitable for genetic manipulation, and we selected the TYK-nu cell line on the basis of the induction of FOXM1 upon co-culture with the TME (Suppl. Fig. 3C). Lentiviral-mediated shRNA transduction was applied to assess the impact of ablating FOXM1 (Suppl. Fig. 3D) on stem-related traits both *in vitro* and *in vivo*. As shown in Fig. 2D, the knockdown of FOXM1 reduced the ability of TYK-nu cells to form spheres.

Tumor initiation is a key defining property of CSC; we therefore tested the effect of FOXM1 ablation on the tumor-initiating ability of OC cells. To this aim, an *in vivo* extreme limiting dilution assay (ELDA) ^20^ was performed with TYK-nu cells knocked down for FOXM1 expression vs control cells. Nude mice were injected subcutaneously with a decreasing number of cells (from 5*10^6 to 1*10^4 cells/mouse) and monitored for tumor formation. ELDA calculation was based on the number of mice with palpable tumors at 21 days after the injection (Suppl. Fig. 3E), which revealed that FOXM1 knockdown reduced the frequency of tumor-initiating cells by 4.6 times (Fig. 2E). Overall, these data demonstrated that FOXM1 is required to sustain stem-like traits and tumor initiation capacity in OC cells.

### TME-dependent induction of FOXM1 is mediated by FAK/YAP signaling

In an attempt to elucidate the molecular mechanisms that underlie FOXM1 induction by TME in OCSC, we first tested the possible involvement of soluble factors. This, however, appeared not to be the case since treatment of OCSC with TME-derived conditioned medium did not induce FOXM1 expression (data not shown). Intercellular contact often stimulates integrin signaling ^21^, which is known to induce FOXM1 expression ^22^. Therefore, we investigated the potential role of Focal Adhesion Kinase (FAK), a classical effector of the integrin pathway ^21^, in TME-dependent upregulation of FOXM1. FAK was not active in TYK-nu OCSC when cultured alone as spheres, where also FOXM1 was not expressed (Fig. 3A). In contrast, when TYK-nu sphere-derived cells were cultured on the TME, FAK was strongly activated concomitantly with the induction of FOXM1 (Fig. 3B).

**Figure 3.**
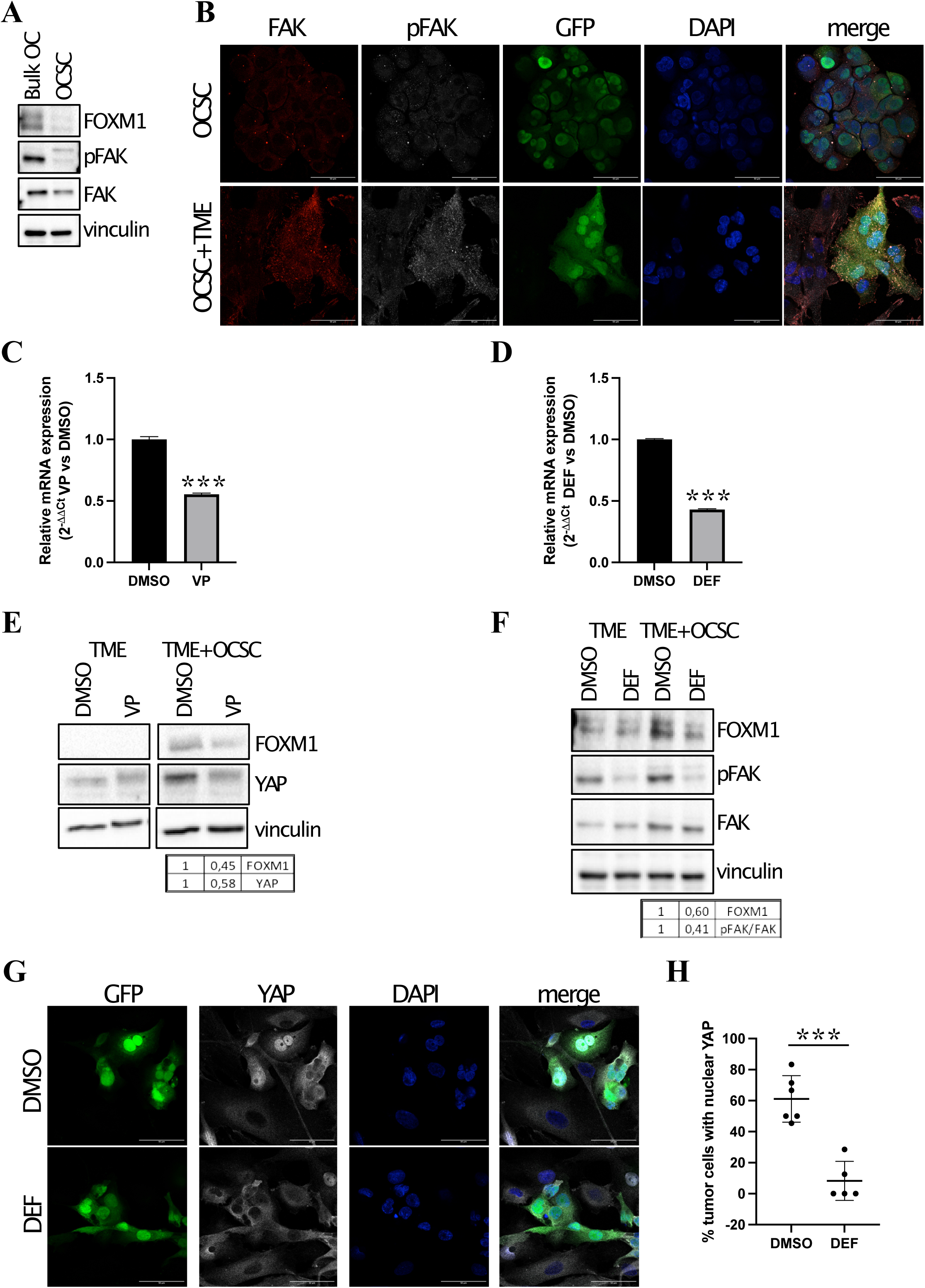
Contact with TME induces FOXM1 signaling in OCSC through FAK/YAP signaling. (A) Cell lysates from TYK-nu Bulk OC and OCSC were immunoblotted for FOXM1, pFAK and FAK, while vinculin was used as loading control. (B) Immunofluorescence for FAK (in red) and pFAK (in grey) on GFP+ TYK-nu OCSC (in green) alone (upper panel) or co-cultured with TME (lower panel). Nuclei were counterstained with DAPI (blue). Scale bar: 50 μm. (C) mRNA expression of *FOXM1* was analyzed by qRT-PCR in TYK-nu OCSC cultured with TME for 24h and treated with vehicle (DMSO, black) or with Verteporfin 3 μM (VP, grey). Data are represented as a relative mRNA expression (2^-1ΔΔCt^), compared to OCSC treated with DMSO. Comparisons between experimental groups were done with two-sided Student’s t-test; \*\*\**p* < 0.005. (D) mRNA expression of *FOXM1* was analyzed by qRT-PCR in TYK-nu OCSC cultured with TME for 24h and treated with vehicle (DMSO, black) or with Defactinib 1 μM (DEF, grey). Data are represented as a relative mRNA expression (2^-1ΔΔCt^), compared to OCSC treated with DMSO. Comparisons between experimental groups were done with two-sided Student’s t-test; \*\*\**p* < 0.005. (E) Cell lysates from TME cultured or not with TYK-nu OCSC and treated for 24h with vehicle (DMSO) or Verteporfin 3 μM (VP) were immunoblotted for FOXM1 and YAP, while vinculin was used as loading control. The panel shows a single blot, intervening lanes were removed for clarity reasons. (F) Cell lysates from TME cultured or not with TYK-nu OCSC and treated for 24h with vehicle (DMSO) or Defactinib 1 μM (DEF) were immunoblotted for FOXM1, pFAK and FAK, while vinculin was used as loading control. (G) Immunofluorescence for YAP (in grey) on GFP+ TYK-nu OCSC (in green) co-cultured with TME for 24h and treated with vehicle (DMSO) or with Defactinib 1 μM (DEF). Nuclei were counterstained with DAPI (blue). Scale bar: 50 μm. (H) Quantification of the experiment shown in panel E. Each dot represents the percentage of OCSC positive for nuclear YAP signaling in a field. The graph shows mean±SD of at least 5 fields. Comparisons between experimental groups were done with two-sided Student’s t-test; \*\*\**p* < 0.005.

One of the pathways stimulated by FAK is YAP signaling ^23^, and FOXM1 is a direct transcriptional target of YAP ^24,25^. Thus, YAP signaling could be activated by TME in OCSC downstream to FAK activation, resulting in FOXM1 upregulation. In support of this hypothesis, GSEA showed significant enrichment for YAP signature not only in OCSC upon co-culture with TME but also in high-FOXM1 vs. low-FOXM1 samples in the TCGA-OC dataset (Suppl. Fig. 4A). On this basis, we investigated the role of a FAK/YAP axis in the regulation of FOXM1. TYK-nu OCSC were cultured on TME in the presence of the YAP inhibitor Verteporfin or the FAK inhibitor Defactinib. Both compounds reduced the TME-mediated induction of FOXM1 (Fig. 3C-F). These data implicated both FAK and YAP in FOXM1 regulation in the OCSC/TME co-culture setting. We could then demonstrate that, on one hand, TYK-nu cells treated with Defactinib displayed a reduction in YAP nuclear localization (Suppl. Fig. 4B) and, on the other, Defactinib significantly reduced YAP nuclear localization in TYK-nu OCSC cultured on TME (Fig. 3G, H). Thus, in our cellular model FAK and YAP act in the same axis.

Taken together, this set of data describes a novel cascade whereby peritoneal TME induces the activation of a FAK-YAP axis in OCSC that ultimately leads to FOXM1 induction

### Interfering with FOXM1 pathway impairs the survival of OCSC on TME and synergizes with PARP inhibitors

The TME-induced FOXM1 activation may represent a specific vulnerability of OCSC and, therefore, interfering with this process could unveil a promising stem cell-eradicating approach. To verify this hypothesis, we first tested if the pharmacological inhibition or expression downmodulation of FOXM1 could affect TME-mediated activation of its downstream pathway. The expression of *FOXM1* itself and of its classical target genes (i.e., *AURKA*, *CCNB1*, *CDK1*, *PLK1*) upon co-culture with TME was analyzed in primary OCSC from HGSOC1 patient in the presence or absence of Thiostrepton, as well as in FOXM1-knockdown vs control OCSC from TYK-nu cells. Both FOXM1 inhibition (Fig. 4A) and its silencing (Fig. 4B) reduced significantly the TME-dependent upregulation of FOXM1 target genes. Moreover, primary OCSC re-isolated from co-cultures treated with Thiostrepton exhibited a significant decrease in sphere formation than OCSC from untreated co-cultures (Fig. 4C). Overall, these data suggested that blocking the TME-mediated induction of FOXM1 might be a strategy to interfere with the stem-related properties of OCSC.

**Figure 4.**
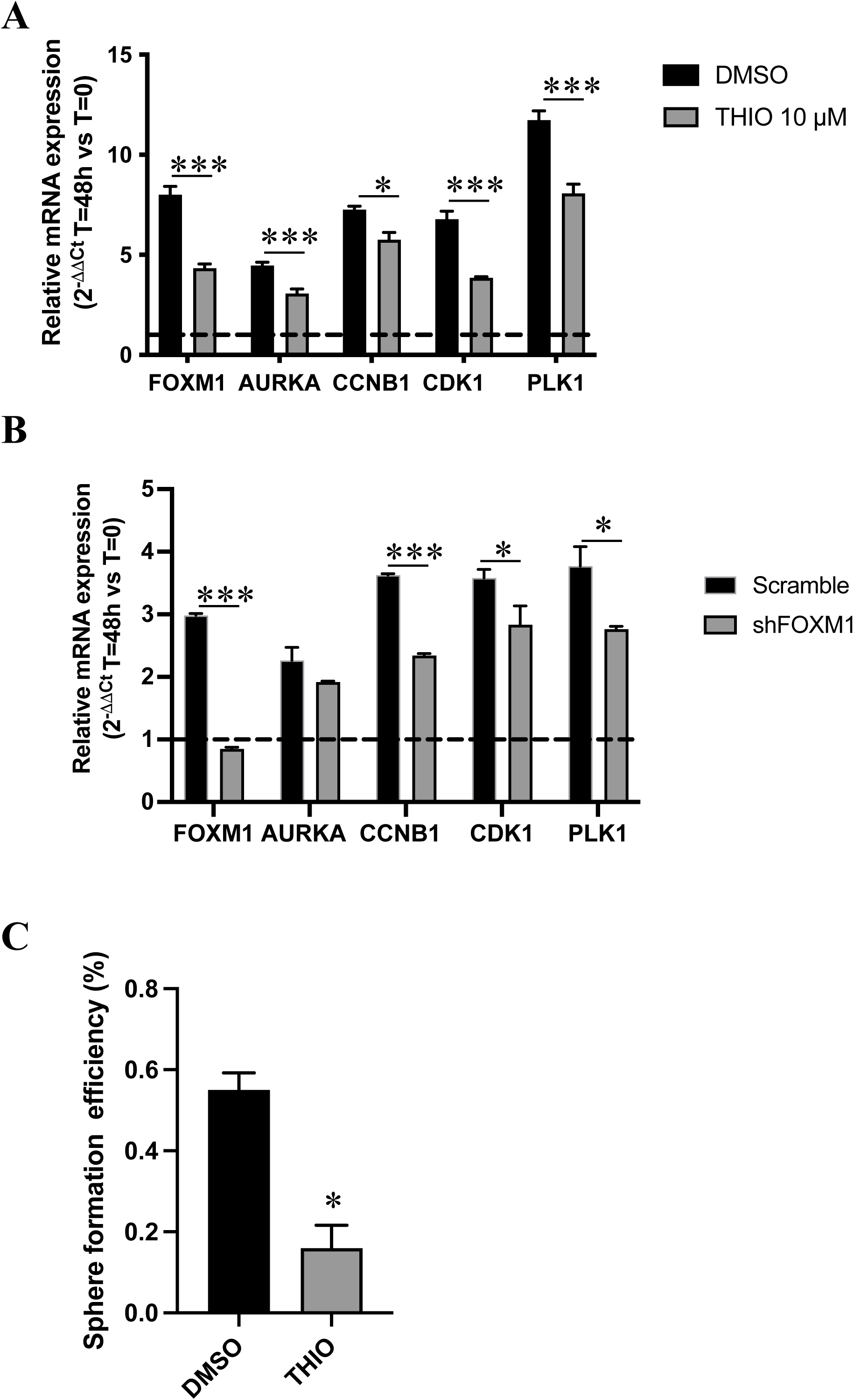
TME stimulates the activation of FOXM1 pathway in OCSC. (A) mRNA expression of *FOXM1* and of its target genes (*AURKA*, *CCNB1*, *CDK1* and *PLK1*) was analyzed by qRT-PCR in HGSOC1 OCSC cultured with TME for 48h and treated with vehicle (DMSO, black) or with Thiostrepton 10 μM (grey). Data are represented as a relative mRNA expression (2^-1ΔΔCt^), compared to cells grown in absence of TME (dashed line). Comparisons between experimental groups were done with two-sided Student’s t-test; \**p* < 0.05, \*\*\**p* < 0.005. (B) mRNA expression of *FOXM1* and of its target genes (*AURKA*, *CCNB1*, *CDK1* and *PLK1*) was analyzed by qRT-PCR in TYK-nu Scramble (black) or shFOXM1 (grey) OCSC cultured with TME for 48h. Data are represented as a relative mRNA expression (2^-1ΔΔCt^), compared to Scramble OCSC not cultured with TME (dashed line). Comparisons between experimental groups were done with two-sided Student’s t-test; \**p* < 0.05, \*\*\**p* < 0.005. (C) HGSOC1 OCSC FACS-sorted after co-culture with TME in presence or absence of Thiostrepton 10 μM (experiment shown in panel A), were subjected to a sphere formation assay in absence of further treatments. Comparisons between experimental groups were done with two-sided Student’s t-test; \**p* < 0.05.

To explore such a possibility, we first generated TYK-nu cells stably tagged with red fluorescent protein (RFP), which were then cultured either as bulk, adherent cells or as OCSC on the TME for 6 days in the presence or absence of Thiostrepton. While Thiostrepton did not affect the survival of adherent cells, it significantly reduced the viability of OCSC (Fig. 5A, B), thus suggesting that FOXM1 inhibition could target the stem cell compartment of HGSOC in the context of the omental niche.

**Figure 5.**
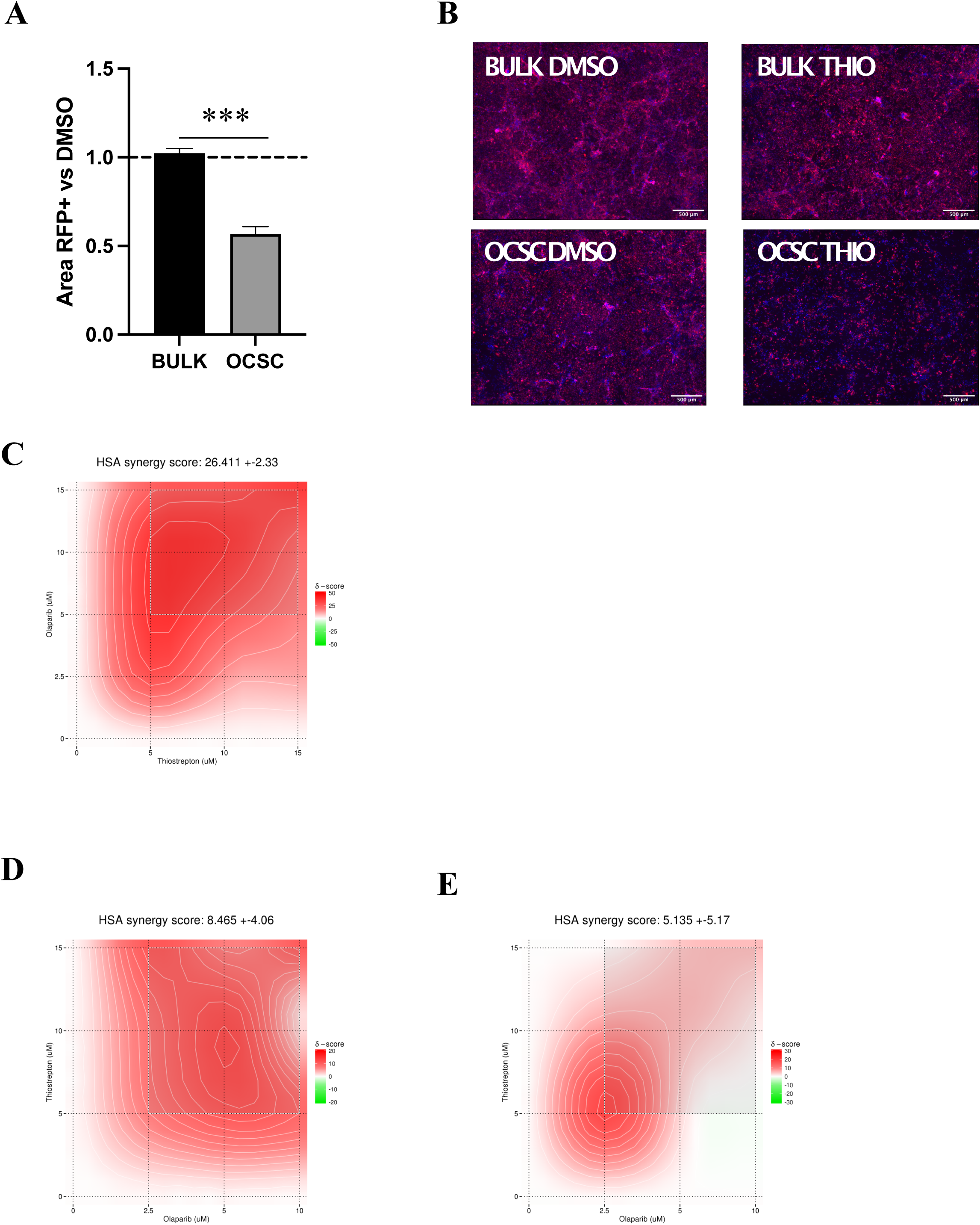
FOXM1 inhibition decreases survival of OCSC cultured on TME and synergizes with PARP inhibitors. (A) RFP positive TYK-nu bulk and OCSC were cultured on TME in presence of Thiostrepton 10 μM, and RFP positive area was measured after 6 days of treatment. Data are expressed as mean RFP^+^ area ± SEM from 3 independent experiments. Comparisons between experimental groups were done with two-sided Student’s t-test; \*\*\**p* < 0.005. (B) Representative pictures from one of the experiments analyzed in panel A. Scale bar: 500 μm. (C) RFP positive TYK-nu OCSC were cultured on TME and treated with 3 different doses of Thiostrepton (5, 10 and 15 μM) and 4 doses of Olaparib (2.5, 5, 10 and 15 μM), alone or in combination. RFP positive area was measured after 6 days of treatment. The combinatorial matrix shows the synergy distribution for the drug combinations according to the Gaddum’s non-interaction (HSA) model, calculated using the SynergyFinder Web Application. The intensity of red color reflects the strength of synergism of each drug concentration. The panel shows the result of one representative experiment (n=3). (D, E) GFP positive OCSC from HGSOC9 (D) and HGSOC10 (E) primary sample were cultured on TME and treated with 3 different doses of Thiostrepton (5, 10 and 15 μM) and 3 doses of Olaparib (2.5, 5 and 10 μM), alone or in combination. GFP positive area was measured after 6 days of treatment. The combinatorial matrix shows the synergism distribution for the drug combinations according to the Gaddum’s non-interaction (HSA) model, calculated using the SynergyFinder Web Application.

Based on these results, the ability of FOXM1 inhibition to target OCSC would be particularly relevant in the maintenance therapy setting, which aims at preventing or delaying OCSC-driven tumor recurrence once the bulk of the tumor has been removed by surgery and cytotoxic treatments. PARP inhibitors (PARPi) are commonly used as maintenance therapy in HGSOC patients with homologous recombination defects ^26–28^; since in other cancer types FOXM1 inhibitors synergized with this class of drugs ^29,30^, we decided to investigate the effect of co-inhibiting PARP and FOXM1 in OCSC co-cultured with the TME. TYK-nu/RFP sphere-derived cells were treated for 6 days with different doses of the PARPi Olaparib and Thiostrepton, alone or in combination. We observed a strong synergism between PARPi and FOXM1 inhibition in reducing the number of residual TYK-nu OCSC cultured on TME (Fig. 5C). To validate these findings in a more clinically relevant setting, we performed the same experiment on OCSC derived from two distinct HGSOC primary samples, and in both we observed a trend towards a synergistic, or at least additive, effect (Fig. 5D, E). Overall, these data demonstrated that the TME-induced upregulation of FOXM1 enhances the sensitivity of OCSC to PARPi.

## Discussion

The TME has long been known to influence the pathophysiology of tumor cells, affecting all phases of cancer progression as well as its responsiveness to treatments ^6^. Particularly intriguing, in this regard, is the impact of TME on CSC: given the contribution of CSC to tumor initiation, dissemination, recurrence and drug resistance ^4^, dissecting the regulatory circuits that link the TME to their function may unveil vulnerabilities that could lead to tumor-eradicating therapies.

In the context of HGSOC, the peritoneum is the primary site of OCSC-driven dissemination and recurrence ^31^. Thus, its crosstalk with OCSC is expected to play a central role in tumor malignancy and to have relevant clinical implications. To investigate the TME/OCSC interplay in a clinically relevant setting, we have set up a platform of organotypic co-culture models, entirely built on patient-derived primary cells, which mimic the 3D architecture of the omental surface ^9,10^. Based on the architecture of the model and on the use of ascites-derived tumor cells, our approach recapitulates to a certain extent the initial contact between metastatic HGSOC cells present in the ascites and the omental niche during early peritoneal dissemination.

The first observation that derived from a global, unbiased transcriptome profiling of patient-derived organotypic co-cultures was that OCSC respond to the contact with peritoneal TME with the activation of pathways related to cell cycle, which is not observed in bulk cancer cells. This may be consistent with a pro-survival function of the TME which may underlie a (transient?) expansion of the OCSC pool and/or may trigger the initiation of the metastatic lesion of HGSOC.

In the transcriptome analysis, co-culture of OCSC with TME induced FOXM1. Previously, this transcription factor was identified as part of a major network involved in the pathophysiology of HGSOC in the integrated multi-omics analysis of The Cancer Genome Atlas (TCGA) project ^12^. Those findings, which support the ability of our organotypic approach to unveil pathways of clinical relevance, were subsequently confirmed and extended in several studies that implicated FOXM1 in HGSOC progression ^13^. Nevertheless, our knowledge on the functional contribution of FOXM1 to OC stemness is scattered and derives from studies performed on cell lines with limited disease relevance ^16,18^. Furthermore, FOXM1 has never been analyzed in the context of OCSC/TME interactions. A previous study on cancer cell lines, in fact, has reported the role of FOXM1 on OC cell adhesion to mesothelium, yet without investigating specifically the CSC subpopulation ^22^. By capitalizing on a fully patient-derived model, our study provides the first evidence of FOXM1 as a pivotal effector of TME-induced stemness features in HGSOC cells. As a corollary to our findings, such a mechanism may contribute to the regulatory role of the peritoneal niche on OCSC-driven dissemination, relapse and chemoresistance of HGSOC. In support of this paradigm, FOXM1 inhibition reduced peritoneal seeding in a syngeneic mouse model of OC ^22^ and sensitized OC cell lines to cisplatin and paclitaxel ^32^.

On the mechanistic level, organotypic co-cultures revealed that the induction of FOXM1 required the adhesion of OCSC to peritoneal TME and was mediated by FAK signaling. This is the first formal demonstration of a causal link between FAK activation and FOXM1 expression. The involvement of FAK in FOXM1-driven OC stemness is relevant given the many clinical trials that are assessing FAK inhibition in different tumor types including OC (https://clinicaltrials.gov/search?cond=ovariancancer&intr=Defactinib). Particularly intriguing, in this context, are the attempts to use FAK inhibition to re-sensitize OC to chemotherapy, which are being pursued also in the clinical setting (https://clinicaltrials.gov/study/NCT03287271) with the support of preclinical data implicating FAK activity in OC chemoresistance ^33,34^. Based on our findings, it is tempting to speculate that the ability of FAK inhibitors to restore chemosensitivity reflects, at least to a certain extent, the disruption of a FAK-FOXM1 axis that underlies OCSC-dependent drug resistance.

Our study clearly points to FOXM1 activity as a vulnerability in the pathophysiology of OCSC, implying that FOXM1-targeted treatments might offer a viable strategy to eradicate OC by eliminating its stem cell compartment. Despite its mechanism of action has not been fully clarified, Thiostrepton (also known as RSO-021) is the most widely used inhibitor of FOXM1, given its well-established ability to counteract this transcription factor, and has been often used in mice showing no major toxicity ^35–37^. Until recently, Thiostrepton has been used only in preclinical studies; however, last year a Phase I/II clinical trial in mesothelioma patients has started (https://clinicaltrials.gov/ct2/show/NCT05278975), which may prelude to the future use of Thiostrepton in the clinical practice and, hence, to its testing in OC patients.

In this context, we showed for the first time the synergism between Thiostrepton and the PARPi Olaparib in reducing OCSC survival on TME. Based on these findings, we argue that the combination of FOXM1 inhibition and PARPi may prove an efficient therapeutic strategy for OC in the maintenance setting, due to its potential to synergistically interfere with OCSC-driven chemoresistance and tumor recurrence.

In summary, we have harnessed clinically relevant, patient-derived organotypic models to profile the impact of peritoneal TME to OCSC at the transcriptional level, which revealed FOXM1 as a novel, TME-induced driver of OCSC pathophysiology. Our data have profound implications not only for a deeper understanding of the pathogenic function of OCSC, but also as a starting point to develop innovative OCSC-eradicating therapies to defeat such a devastating tumor.

## Materials and Methods

### Cell culture

The human ovarian cancer cell line TYK-nu was a kind gift of P. Lo Riso and G. Testa and was grown in MEM medium (Sigma-Aldrich, cat#G6148) containing 10% fetal bovine serum (FBS, Biowest cat# S1860), 2 mM L-glutamine (Euroclone, cat# ECB3000), 100 U/ml penicillin, 100 μg/ml streptomycin (Euroclone, cat# ECB3001). The cell line was routinely tested for mycoplasma with a PCR-based method and authenticated via short tandem repeat profiling.

Primary cell cultures were established from peritoneal ascites of high-grade serous ovarian cancer (HGSOC) patients. Tumor samples were provided by the Division of Gynecology at the European Institute of Oncology (Milan) upon informed consent from patients undergoing surgery. Clinicopathological features of the samples used in this work are provided in Supplementary Table S1. Tumor histology was confirmed by a board-certified pathologist (G. Bertalot), while the identity of cancer cells was confirmed by immunostaining for cytokeratins 5, 7, and 8, or for pan-cytokeratins. The purity of primary cell culture was consistently over 95%. Tissue isolation and culture conditions of primary cells were performed as described previously ^38^.

Mesothelial cells and fibroblasts were isolated from healthy omental specimens. To isolate mesothelial cells, the omental tissue was washed several times with sterile PBS, which was then collected and centrifuged at 1400 rpm for 5 minutes. Cell pellet containing mesothelial cells was cultured with RPMI 1640 (Euroclone, cat# ECM2001L), 20% FBS, 2 mM L-glutamine, 100 U/ml penicillin, 100 μg/ml streptomycin, 1% NEAA (Lonza, cat# 13-114E), 1% MEM vitamins (Lonza, cat# 13607C). To isolate fibroblasts, the tissue was minced and incubated overnight on an orbital shaker with 100 U of hyaluronidase (Merck, cat# H3884) and 1,000 U of collagenase type III (Worthington Biochemical, cat# LS004183) in DMEM (Euroclone, cat# ECM0103L), 10% FBS at 37°C. The digested tissue was then centrifuged at 1400 rpm for 5 minutes, the cell pellet containing fibroblasts was washed with phosphate-buffered saline (PBS) and cultured in DMEM, 10% FBS, 1% glutamine, 100 U/ml penicillin, 100 μg/ml streptomycin, 1% NEAA, 1% MEM vitamins. Mesothelial cells and fibroblast were used for the experiments within three passages.

All cell lines and primary samples were maintained at 37°C in a humidified incubator with 5% CO_2_. When needed, cells were treated with the following reagents: Thiostrepton and Verteporfin from Sigma-Aldrich (cat# 598226 and SML0534, respectively); Defactinib and Olaparib from Selleckchem (cat# S7654 and S1060, respectively).

### Sphere Formation Assay

Sphere formation assays were performed as described ^39^. Briefly, single cell suspensions, derived from ovarian cancer cell lines or primary samples, were seeded at low density under non-adherent conditions in poly-(2-hydroxyethyl methacrylate) coated dishes (Sigma-Aldrich, cat# P3932) and allowed to form clonal spheres. TYK-nu cells were seeded at a density of 5000 cells/ml, and OCSC-enriched sphere were maintained in DMEM:F-12 (1:1) (Gibco, cat# 11320033), supplemented with 2% B27 (Thermo Fisher Scientific, cat# 17504044;), 2 mM L-glutamine, 100 U/ml penicillin, 100 μg/ml streptomycin, 20 ng/mL epidermal growth factor (EGF, Merck, cat# E4127), and 10 ng/mL fibroblast growth factor-2 (FGF2; Peprotech, cat# AF-100-18B). Primary OCSC cultures were seeded at 5000 cells/ml in MEBM (Lonza, cat# CC-3151) supplemented with 2 mM L-glutamine, 100 U/mL penicillin, 100 µg/mL streptomycin, 5 µg/mL insulin (Thermo Fisher Scientific, cat# RP10935), 0.5 µg/mL hydrocortisone (Sigma-Aldrich, cat# H0888), 1 U/mL heparin (Voden, cat # 07980), 2% B27, 20 ng/mL EGF, and 20 ng/mL FGF2.

Spheres were then dissociated with StemPro Accutase (Thermo Fisher Scientific, cat# A1110501), according to the manufacturer’s protocol, and re-plated under the same conditions to obtain second-generation spheres or used for co-cultures with the TME.

Sphere formation was assessed 5–10 days after seeding. The sphere-forming efficiency (SFE) was defined as the ratio between the number of spheres counted and the number of cells seeded.

### Cell viability

Ovarian cancer cells were seeded in 96-well plates in quadruplicates at a density of 4000 cells/well, and after 24 hours they were treated with different doses of Thiostrepton. After 72 hours of treatment, the metabolic activity was quantified using the Cell counting kit-8 (Sigma-Aldrich, cat# 96992), following the manufacturer’s instructions. The absorbance at a wavelength of 450 nm was measured using the Glomax Plate Reader.

### 3D organotypic model

The 3D organotypic model was assembled as previously described ^14^. Briefly, primary omental fibroblasts were resuspended in a medium containing 5 μg/ml collagen-I (Corning, cat# 354236) and 5 μg/ml fibronectin (Corning, cat# 354008), and then plated and incubated at 37°C. After 4 hours, the fibroblast layer was overlaid with primary mesothelial cells and the co-culture was incubated at 37°C for 2 days. Single-cell suspensions of OC cell lines or of primary HGSOC cells, pre-labeled with 0.1 µM CMFDA (Thermo Fisher Scientific, cat# C2925) according to manufacturer’s instructions, were seeded on top of 3D organotypic cultures. After 48 hours, the co-cultures were dissociated to single cells and CMFDA-labeled HGSOC cells were isolated by fluorescence-activated cell sorting (FACS) and subjected to RNA extraction.

In some experiments, the 3D organotypic model was assembled in Nunc Lab-Tek chamber slides for immunofluorescence analysis, or in 96-well plates to assess response to drug treatments. In the latter case, GFP or RFP positive HGSOC cells were seeded on the TME, and after 24 hours different drugs or drug combinations were added. Each treatment was performed in triplicate. After 5 days of treatment, nuclei were counterstained with Hoechst 33342 (Sigma-Aldrich, cat# 2261), and the fluorescent signal coming from GFP (or RFP) and Hoechst was acquired using the Leica THUNDER imaging system. Nine fields per well were acquired, and the area positive for GFP (or RFP) and Hoechst was quantified using the Fiji software. The GFP+ (or RFP+) area of treated cells was normalized on the one of vehicle-treated controls, to obtain the percentage of cells that survived at the end of treatment. For the drug combinations, the SinergyFinder web-tool (https://synergyfinder.fimm.fi/; ^40^) was then used to quantify the degree of synergy, or antagonisms, of the tested combinations, using the Highest Single Agent (HSA) synergy scoring model. The resulting synergy scores were visualized as a 2D synergy map, which highlighted in red the drug concentrations displaying a synergistic effect, in green the ones with an antagonistic effect.

### RNA extraction and qRT-PCR

Total RNA was extracted using the RNeasy Mini Kit (QIAGEN, cat# 217004) according to the manufacturer’s protocol and quantified using the Nanodrop instrument (Thermo Fisher Scientific). RNA quality was checked using an Agilent 2100 Bioanalyzer. Preparation of cDNA and qRT-PCR were performed by the Cogentech qPCR Service (Milan, Italy).

Gene expression levels for *FOXM1* and its target genes were analyzed and normalized against the housekeeping genes *GAPDH* and *HPRT1*. TaqMan assays for specific genes are listed in Supplementary Table S5. Normalized expression changes were determined with the comparative threshold cycle (2^−ΔΔCT^) method.

### RNA-Seq analysis

Poly-A enriched strand-specific libraries were generated with the TruSeq mRNA V2 sample preparation kit (Illumina, cat# RS-122-2001), ribosomal RNA depleted strand-specific RNA libraries with the TruSeq Stranded Total RNA LT sample preparation kit with Ribo-Zero Gold (Illumina, cat# RS-122-2301 and #RS-122-2302), and transcriptome capture based libraries with the TruSeq RNA Access Library Prep Kit (Illumina, cat# RS-301-2001). All protocols were performed following the manufacturer’s instructions. Libraries were sequenced by Illumina HiSeq2000 resulting in paired 50nt reads. The sequencing coverage and quality statistics for each sample are summarized in Supplementary Table S6.

Fastq files were aligned to the hg38 genome assembly using STAR v 2.7.5c (PMID: 23104886). STAR gene counts were filtered by selecting those genes with raw counts > 10 in at least 2 out of 6 samples per each group of the comparison. Filtered gene counts were then normalized by the median of ratios method implemented in DESeq2 R package (PMID: 25516281). DESeq2 R package was used also to select differentially expressed genes between -TME (T0) and +TME (T48) in OCSC and bulk OC conditions. Heatmaps were generated by using Cluster 3.0 for Mac OS X (C Clustering Library 1.56) and Java TreeView version 1.1.6r4 (uncentered correlation and centroid linkage) using median centered log_2_ data.

Log_2_FC data were subjected to Core Analysis of Ingenuity Pathway Analysis (IPA; QIAGEN) setting the following parameters: i) genes and endogenous chemical as Reference Set; ii) direct and indirect relationship; iii) human as Species. Canonical Pathways and Upstream Regulators results of the Core Analysis were considered.

Gene Set Enrichment Analysis (GSEA; https://www.gsea-msigdb.org/gsea/index.jsp; PMID: 12808457 and 16199517) was performed using the “xtools.gsea.GseaPreranked” tool to input expression data ordered by the “Stat” output of DESeq2 R package (i.e., Wald statistic) in the case of TME in OCSC condition. In the case of TCGA-OC dataset, GSEA was performed by correlating expression data to FOXM1 expression profile by Pearson correlation metric. One-thousand random gene sets permutation was performed for false discovery rate (FDR) computation of the normalized enrichment scores (NES). Hallmark gene sets (N=50) (https://www.gsea-msigdb.org/gsea/msigdb/human/collections.jsp) or specific gene sets representative of YAP signaling (CORDENONSI_YAP_CONSERVED_SIGNATURE.v2022.1.Hs.gmt) and FOXM1 transcription factor network (PID_FOXM1_PATHWAY.v7.5.1.gmt.txt) were used.

### Immunoblotting

Cells were lysed in hot lysis buffer (2.5% SDS, 125 mM Tris-HCl [pH 6.8]), after 15 minutes incubation at 95°C.

For cytosolic/nuclear fractionation, cells were washed with ice-cold PBS and incubated with a hypotonic buffer (10 mM Hepes pH 7.9, 10 mM KCl, 0.1 mM EDTA, 0.1 mM EGTA, 0.6% NP-40, 1 mM NaVO_3_, 1:500 Protease Inhibitors Cocktail from Sigma-Aldrich cat# P8340) for 20 minutes on ice in a shaker. Cells were then scraped and collected in an eppendorf tube. After centrifugation for 10 minutes at maximum speed, the supernatant containing the cytoplasmic fraction was collected. For nuclear extracts, the pellet was washed 3 times with the same buffer and then incubated with the nuclear lysis buffer (20 mM Hepes pH 7.9, 400 mM NaCl, 0.1 mM EDTA, 0.1 mM EGTA, 50% Glycerol, 1 mM NaVO_3_, 1:500 Protease Inhibitors Cocktail) for 1 hour in a shaker at 4°C. After centrifugation for 5 minutes at maximum speed, the supernatant containing the nuclear fraction was collected.

Protein concentration was determined using a Pierce BCA Protein Assay kit (Thermo Fisher Scientific, Inc, cat# 23227) according to the manufacturer’s instructions. Equal amounts of protein extracts (20 µg) were resolved in acrylamide gel and transferred onto nitrocellulose membranes. The membranes were incubated overnight at 4°C with the following primary antibodies: FOXM1 (Cell Signaling, cat# 5436), FAK (Invitrogen, cat# 39-6500), phospho-FAK (Tyr397) (Invitrogen, cat# 700255), YAP (Cell Signaling, cat# 14074), beta-Tubulin (Santa Cruz, cat# sc-58886), vinculin (Sigma-Aldrich, cat# V9131), Lamin A/C (Santa Cruz, cat# sc-7292). Membranes were incubated with IgG HRP-conjugated secondary antibody (Bio-Rad Laboratories) for 1 hour at room temperature. The signal was detected by the Clarity Western ECL Substrate (Bio-Rad, cat# 1705062) as described in the manufacturers protocol and the images were acquired using ChemiDoc (Bio-Rad) and analyzed with the Fiji software.

### Immunofluorescence

Cultured cells (or cytospins with OCSC-enriched spheres) were fixed with 4% paraformaldehyde for 5 minutes at room temperature and then permeabilized in ice-cold PBS, 0.5% Triton X-100 for 3 minutes at 4°C. After blocking for 1 hour at room temperature with blocking buffer (PBS, 0.2% BSA, 1% donkey serum, 0.05% Tween-20 and 0.02% NaN3), cells were incubated for 2 hours with the following primary antibodies, diluted in blocking buffer: FOXM1 (Cell Signaling, cat# 5436), FAK (Invitrogen, cat# 39-6500), phospho-FAK (Tyr397) (Invitrogen, cat# 700255), YAP (Cell Signaling, cat# 14074). Cells were then washed with PBS and incubated with the Alexa Fluor-conjugated secondary antibodies (Jackson Laboratories) for 1 hour at room temperature. Nuclei were counterstained with DAPI (Sigma-Aldrich, cat# 32670). Images were acquired using the Leica SP8 Confocal microscope.

### Immunohistochemistry

Immunohistochemical (IHC) staining from human tissues was performed on 3-μm-thick sections cut from formalin-fixed paraffin-embedded samples and dried in a 37°C oven overnight.

IHC staining for FOXM1 was performed using Bond III IHC auto-stainer for full Automated Immunohistochemistry (Leica Biosystems). Antigen was unmasked with Tris-EDTA pH 9 (Bond Epitope Retrieval Solution 2, Leica Biosystems, cat# AR9640).

The anti-FOXM1 antibody (Abcam, cat# ab207298) was diluted 1:250 in Bond Primary Antibody Diluent (Leica Biosystems, cat# AR9352), and BOND IHC Polymer Detection Kit (Leica Biosystems, cat# DS9800) was used to stain the antibody with DAB Cromogen. IHC samples were counterstained using Hematoxylin solution (Leica Biosystems, cat# RE7107-CE). Pictures of stained sections were acquired with the Aperio ScanScope XT instrument.

### Lentivirus production and cell transduction

Lentiviral vectors were generated by transient co-transfection of the packaging cell line HEK293T, purchased from ATCC and cultured as described previously ^38^, with 10 μg of the following anti-FOXM1 shRNA from Genecopoeia: sh-C (5’-TAATACGACTCACTATAGGG-3’; HSH096566-LVRU6GP-c); sh-E (5’-TAATACGACTCACTATAGGG-3’; HSH096566-LVRU6GP-e) or with the scrambled control (CSHCTR001-LVRU6GP, Genecopoeia), and the following packaging vectors: PMD2G (3 μg), Rre (5 μg), and REV (2.5 μg), using the calcium phosphate precipitation method. The supernatant from HEK293T, containing the virus particles, was supplemented with 8 μg/ml of polybrene and used to transduce recipient cells.

### *In vivo* limiting dilution assays

Mice were housed under specific pathogen-free conditions in isolated vented cages and allowed access to food and water ad libitum. Six-eight weeks-old nude female mice (from Charles River Laboratories) were injected subcutaneously into the flank with serial dilutions of TYK-nu shSCR and shFOXM1 cells, ranging between 5*10^6^ and 1*10^4^ cells/site in a 1:1 (vol:vol) mixture with growth factor-reduced Matrigel (Corning, cat# 356231) and PBS, with a final volume of 100 µl per injection. Each experimental group consisted of 7 mice.

Tumor latency was defined as the time interval from the injection to the formation of palpable tumors. Tumor take frequency was determined as the number of mice with palpable tumors. Cancer stem cell frequency was measured using the ELDA online software http://bioinf.wehi.edu.au/software/elda.

Body weight and general physical status were monitored daily, and the mice were sacrificed when the tumor reached a volume of 1000 mm^3^.

All experimental procedures involving mice and their care were performed following protocols approved by the fully authorized animal facility of our Institutions and by the Italian Ministry of Health (as required by the Italian Law) (protocol no. 239/2023-PR) and in accordance with EU directive 2010/63.

### Data Availability Statement

RNAseq data are available by request and will be uploaded to public databases prior to publication. The TCGA Ovarian Serous Cystadenocarcinoma (OV) RMA normalized dataset (samples with mRNA data, U133 microarray data only, N=535) was downloaded from GDAC firehose through cBioPortal data portal (https://www.cbioportal.org/study/summary?id=ov_tcga).

### Statistical Analysis

Independent experiments were considered as biological replicates. For *in vivo* experiments, each mouse represented one biological replicate. Data are expressed as means ± SEM, calculated from at least three independent experiments. Statistical significance was evaluated with Student’s two-tailed t test (GraphPad Prism 8). Cut-off threshold to define significance was set at p < 0.05. Asterisks correspond to the p-value calculated by a two-tailed, unpaired t-test (*p < 0.05, **p < 0.01, ***p < 0.001).

## Acknowledgements

We are grateful to all patients who donated their samples for this work, to the Ovarian Cancer Center staff at IEO (Milano) for providing the tumor samples, and to the IEO Biobank staff for collection and annotation of surgical samples. We thank F. Balkwill and B. Malacrida for helpful discussion; G. Bertalot for the histological analysis of tumor specimens; the staff of the Units of Genomics, Molecular Pathology, Cell Culture and FACS at IEO; and P. Lo Riso and G. Testa for kindly providing TYK-nu cell line.

This work was supported by grants from Ovarian Cancer Research Alliance (Collaborative Research Development Grant 648516 to UC, EL), Associazione Italiana Ricerca sul Cancro (IG-21320 to UC; IG-22827 to FB), the Italian Ministry of Health (PE-2016-02362551 to UC; RF-2021-12372433 to FB).

The study funders had no role in the study’s design, collection, analysis, and interpretation of the data, writing of the manuscript, and the decision to submit the manuscript for publication.

Fabrizio Bianchi wishes to dedicate this work to the memory of Lucia, an extraordinary girl who experienced cancer as an opportunity to entrust her and all her loved ones to the love of the Father.

## Notes

### Competing Interest Statement

The authors have declared no competing interest.

