## Supplementary Figures for "Tumor microenvironment-induced FOXM1 regulates ovarian cancer stemness"

**Supplementary Figure 1**.

(A) Gene Set Enrichment Analysis (GSEA) of FOXM1 transcription factor network on the ranked expression data matrix of OCSC after co-culture with TME. NES, normalized enrichment score. FDR, false discovery rate. (B) Immunofluorescence for FOXM1 (in red) on CMFDA-labelled OCSC from HGSOC1 sample (in green) co-cultured with TME, or on HGSOC1 OCSC spheres only (lower panel). Nuclei were counterstained with DAPI (blue). Scale bar: 50 μm.

**Supplementary Figure 2.**

Gene Set Enrichment Analysis (GSEA) of HALLMARK_MITOTIC_SPINDLE, HALLMARK_G2M_CHECKPOINT and HALLMARK_E2F_TARGETS gene sets on ranked expression data matrix of OCSC after co-culture with TME and on the ranked expression data matrix of TCGA-OC cohort correlating with FOXM1 expression profile (High → Low), showing a significant enrichment for all these signatures in both datasets.

**Supplementary Figure 3.**

(A) Cell viability of HGSOC8 primary sample measured after 72h of treatment with vehicle (DMSO) or 3 different doses FOXM1 inhibitor Thiostrepton. (B) Sphere formation efficiency in HGSOC7 and HGSOC8 primary samples treated or not with Thiostrepton 10 μM. Comparisons between experimental groups were done with two-sided Student's t-test; $*p < 0.05$, $***p < 0.005$. (C) mRNA expression of *FOXM1* was analyzed by qRT-PCR in a panel of ovarian cancer cell lines (bulk OC or OCSC) cultured with TME for 48h. Data are represented as a relative mRNA expression ($2^{-\Delta\Delta Ct}$) of bulk (in black) and OCSC (in grey) co-cultured with TME for 48 hours (T=48h) compared to cells grown in absence of TME (T=0, dashed line). Comparisons between experimental groups were done with two-sided Student's t-test; $***p < 0.005$. (D) Cell lysates from TYK-nu cells control (Scramble) and silenced for FOXM1 with 2 different shRNAs (shFOXM1-C and E) were immunoblotted for FOXM1 and beta-tubulin was used as loading control. The panel shows a single blot, intervening lanes were removed for clarity reasons. (E) Nude mice were transplanted subcutaneously with decreasing numbers of either TYK-nu Scramble or TYK-nu shFOXM1-C cells, and analyzed for the presence of a palpable lesion (as a sign of tumor take) 21 days after injection.

**Supplementary Figure 4.**

(A) Gene Set Enrichment Analysis (GSEA) of CORDENONSI_YAP_CONSERVED_SIGNATURE dataset on ranked expression data matrix of OCSC after co-culture with TME and on the ranked expression data matrix of TCGA-OC cohort correlating with FOXM1 expression profile (High →

Low), showing a significant enrichment for YAP signature in both datasets. NES, normalized enrichment score. FDR, false discovery rate (also referred to as q-value). (D) Cytosolic and nuclear lysates from TYK-nu cells treated for 24h with vehicle (DMSO) or Defactinib 1 μM (DEF) were immunoblotted for YAP; beta-tubulin was used as loading control for the cytosolic fractions, while Lamin A/C was used for nuclear fractions. The panel shows a single blot, intervening lanes were removed for clarity reasons.

### Supplementary Figure 1

## A

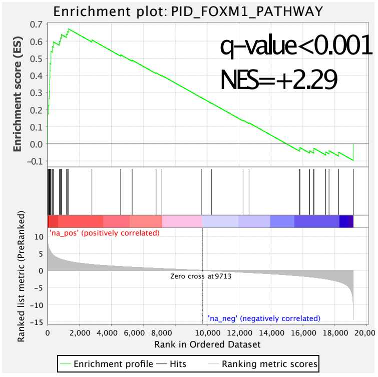


## B

FOXM1/CMFDA/DAPI

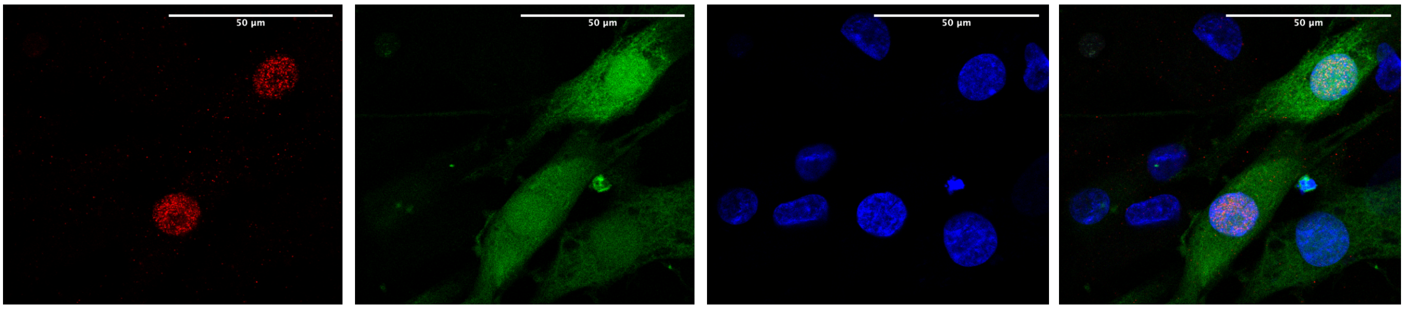


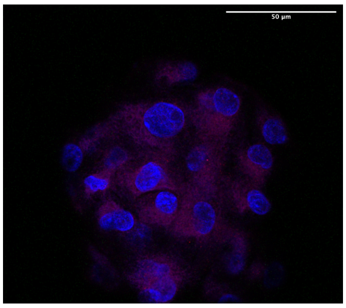

FOXM1/DAPI

### Supplementary Figure 2

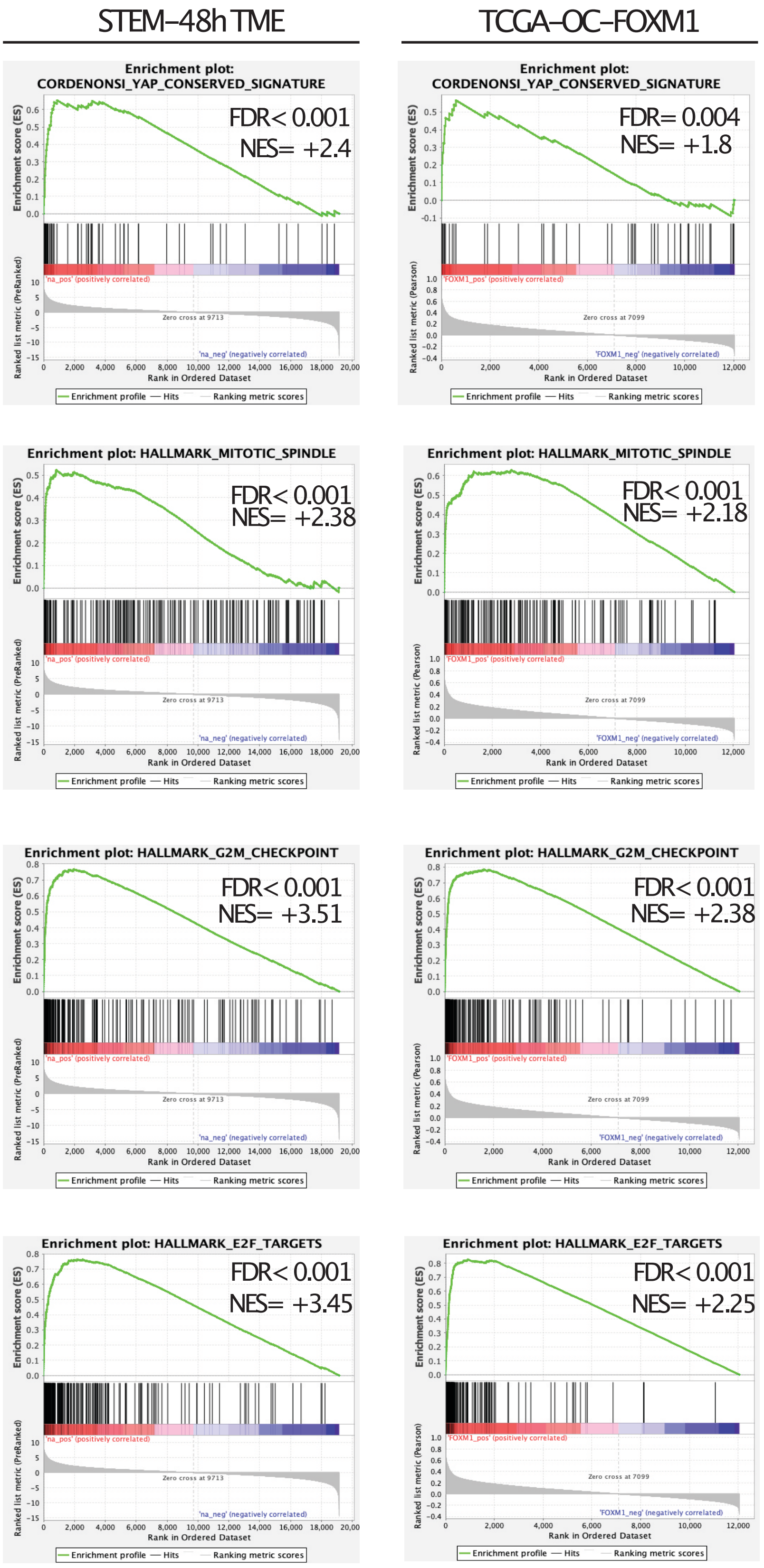

### Supplementary Figure 3

A

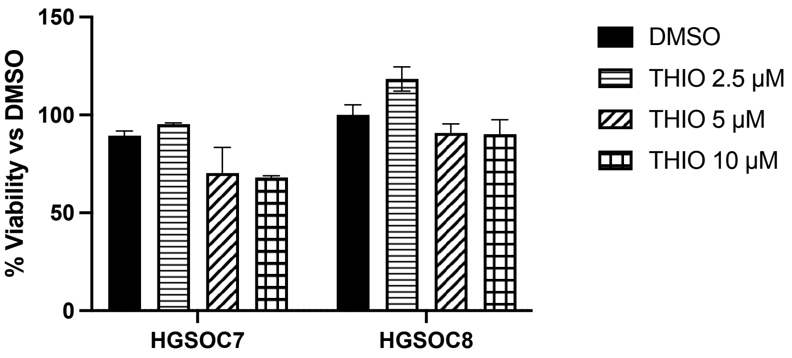


B

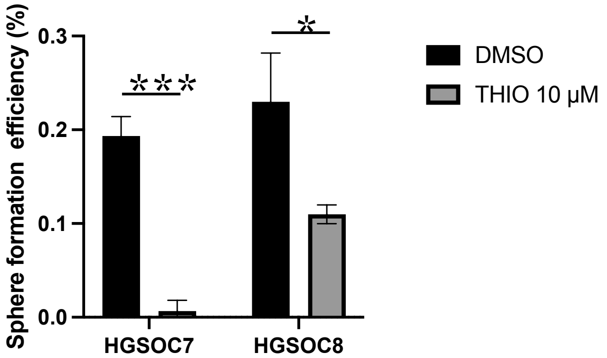


C

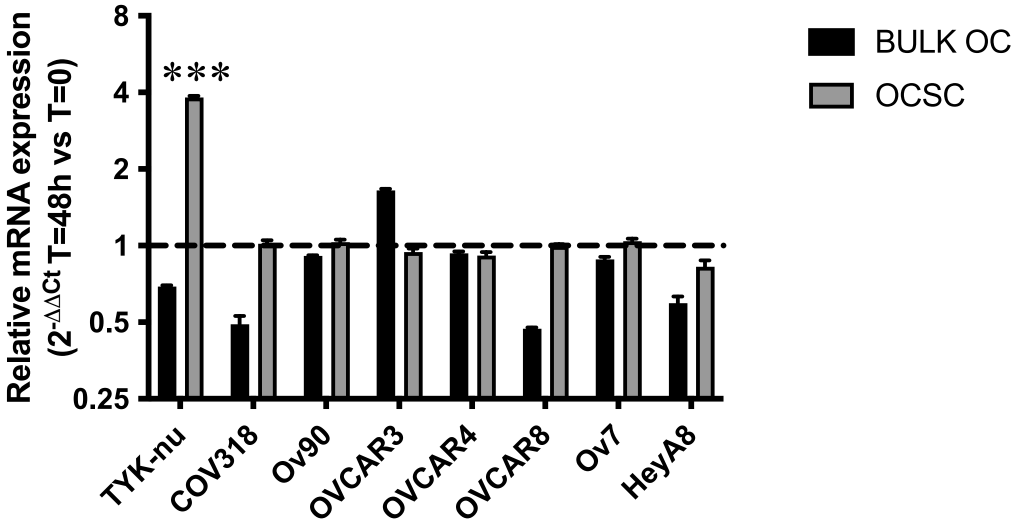


D

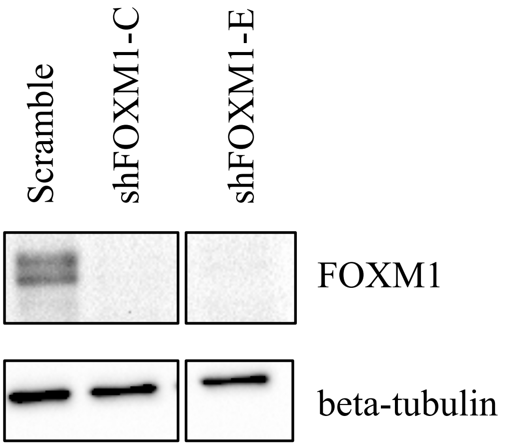


E

| Group | N. of transplanted cells | Tumor take (day 21) |
|---|---|---|
| Scramble | 5*10^6 | 7/7 |
| | 1*10^6 | 7/7 |
| | 2.5*10^5 | 5/7 |
| | 1*10^5 | 6/7 |
| | 1*10^4 | 2/7 |
| shFOXM1 | 5*10^6 | 7/7 |
| | 1*10^6 | 3/7 |
| | 2.5*10^5 | 5/7 |
| | 1*10^5 | 6/7 |
| | 1*10^4 | 0/7 |

### Supplementary Figure 4

## A

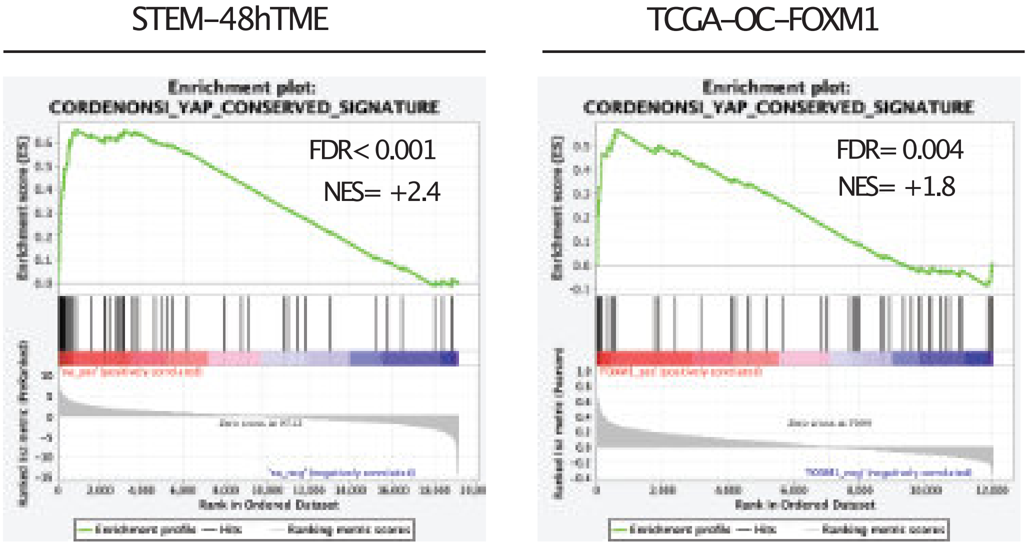


## B

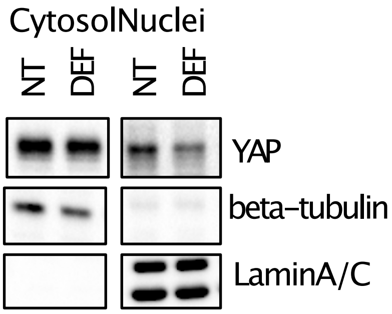
